# Targeting MAPAKAP2(MK2) to combat inflammation by avoiding the differential regulation of anti-inflammatory genes by p38 MAPK inhibitors

**DOI:** 10.1101/2022.07.17.500377

**Authors:** Rakesh Kumar Singh, Reena Sodhi, Sameer Sharma, Sunanda Ghosh Dastidar, Ruchi Tandon

## Abstract

p38 mitogen-activated protein kinase (p38 MAPK) plays an important role in the key cellular processes related to inflammation. Several small molecule inhibitors of p38 MAPK therefore have been evaluated for their anti-inflammatory potential and progressed from early discovery to late phase clinical trials. Most of these efforts however have failed due to severe toxicity concerns. Since p38 MAPK has several downstream substrates, inhibition of p38 MAPK, therefore, leads to the modulation of all its substrates, resulting into a dis-balance of pro- and anti-inflammatory response and multiple toxicity concerns. Targeting p38MAPK MAPKAPK2 (MK2), one of the downstream substrates of p38 MAPK directly, is expected to be a better anti-inflammatory approach without having any toxicity concerns. In this manuscript, we are reporting biological data of representative MK2 inhibitor to validate its anti-inflammatory potential and a comparison of p38 MAPK and MK2 inhibitors in cell based assays to understand their relative toxicities.

## 1. Introduction

Mitogen-activated protein kinase(MAPK) is the key regulator of inflammation. Inflammatory stimulus leads to the activation of three predominant MAPK pathways such as p38 MAPK, extracellular-signal-regulated kinase(ERK) and c-Jun N-terminal kinase(JNK). Out of these, p38 MAPK, has been demonstrated to be the central regulator of inflammation. Inhibiting the p38 MAPK pathway leads to reduced production of pro-inflammatory cytokines, such as TNF-α, IL-6 and IL-1β^1^. Due to its vital role in inflammation, p38 MAPK is therefore an attractive therapeutic target for inflammatory diseases. Several p38 MAPK inhibitors, with diverse chemical structures have been explored till date and shown efficacy in several pre-clinical and clinical disease models including rheumatoid arthritis(RA), Inflammatory bowel disease(IBD), psoriasis, asthma and chronic obstructive pulmonary disease(COPD). None of the p38 MAPK inhibitors, however, have reached to clinic yet. This is most likely due to the unacceptable safety window^2^, transient efficacy profiles or opportunistic infections. The most common side effects associated with p38 MAPK inhibitors are the toxicities related to liver, dermal, gastrointestinal and central nervous systems. There are reports suggesting that one of the possible reasons for the toxicities observed with p38 inhibitors is that they increase the activation of JNK. In addition, p38 MAPK inhibitors also inhibit the phosphorylation of anti-inflammatory mediator Msk1/2^3^ leading to an imbalance of pro- and anti-inflammatory properties.

MAPK being a predominant inflammatory pathway, several members of this signaling network have been explored in addition to p38 MAPK, to identify a potential molecular target with desired anti-inflammatory profile^21,23^. Targeting a molecule downstream to p38 MAPK, such as MAPK-activated protein kinase 2(MK2), seems to be an alternate and safer approach^4-6^. MK2 is a serine/threonine kinase and is activated directly through phosphorylation of p38 MAPK by stress and inflammatory stimuli. Inhibition of MK2 therefore reduces the levels of pro-inflammatory cytokines but spares several other substrates of p38 MAPK and upstream molecules^7^.

Enormous data has emerged to witness the potent anti-inflammatory properties of p-38-MAPK inhibitors^10,11,22^ along with convincing results emerging with regard to the anti-inflammatory properties of MK2 inhibitors as well. Serum levels of TNF-α and IL-6 were found to be significantly low in LPS induced endotoxemia model using MK2 knock-out mice. MK2 knock-out mice were also resistant to collagen induced arthritis^8,9^. These studies provide the *in vivo* proof of concept for MK2 inhibition for ant-inflammatory properties.

In order to dig out the differential toxicities associated with the two approaches, we conducted *in vitro* experiments to understand the mechanistic differences between the two anti-inflammatory approaches at the cellular level using p38 MAPK and MK2 inhibitors. It is interesting to note that although both the molecular targets, belong to the same signaling pathway, consequences of inhibiting the two vary significantly. Our provides the mechanistic insights into the clinical concerns associated with p-38 inhibitors and why inhibiting MK2 can be a better approach. This data can also be seen in the light of the knock out data of p38 MAPK and MK2 in mice. MAPK knock-out mice although are embryonically lethal, MK2 knock-out mice have a normal phenotype.

## 2. Materials and Methods

### 2.1 Materials

#### 2.1.1 Reagents

MK2 enzyme assays were performed using HTRF KinEASE STKS1 kit from Cisbio HTRF technology. Human recombinant MK2 enzyme was procured from Millipore. DuoSet kits were purchased from R& D Systems and cytokines ELISA kits were procured from eBiosciences. Antibodies were purchased from Cell Signaling Technologies. All other chemicals and reagents were procured from Sigma-Aldrich, unless stated otherwise. BIRB-796 and PF-3644022 were procured commercially from Sigma and Tocris respectively. All antibodies were procured from cell signaling technology.

#### 2.1.2. Cells and cell culture

U937 cells were maintained in suspension culture in RPMI-1640 medium, supplemented with 10%(v/v) heat-inactivated fetal bovine serum(FBS), 2 mM L-glutamine and 10 mM HEPES, at 37°C in a humidified atmosphere of 5% CO_2_.

Blood was freshly drawn from healthy volunteers after informed written consent in heparinized containers after receiving the informed written consent according to protocol approved by Institutional Ethics Committee(IEC). Isolation of PBMCs from human blood was performed by density gradient centrifugation on Histopaque-1077(Sigma). Briefly, fresh blood was mixed at a ratio of 1:1 with RPMI-1640 media at room temperature and this mixture was layered over Histopaque-1077 in a ratio of 2:1. Tubes were then centrifuged at 1180□x g for 30□min at 20°C; buffy coats were collected, re-suspended in complete RPMI-1640 media and centrifuged at 1180□x g for 10□min at 20°C. Cell pellets were re-suspended in complete RPMI-1640 media and PBMCs were counted.

#### 2.1.3. Animals

Experiments were conducted on healthy in-bred Balb-c mice were obtained from the Institutional Experimental Animal Facility. Animals were housed in standard cages and maintained at a temperature of 24±2 °C with controlled illumination to provide a light–dark cycle of 12 h. All experimental protocols were approved by the Institutional Animal Ethics Committee (IAEC) and experiments were conducted as per the recommendations received from the Committee for the Purpose of Control and Supervision on Experiments on Animals (CPCSEA) Guidelines for the use and care of experimental animals.

### 2.2 Methods

#### 2.2.1. Cell Proliferation assay using MTT

Since several p38 inhibitors have been reported to have liver toxicities, we chose Hep-G2 cell line to evaluate if the reported human toxicities also reflect in the conventional cell proliferation assays using MTT. To test this hypothesis we evaluated p38 MAPK inhibitors SB-203580, BIRB-796 and MK2 inhibitor PF-3644022 in the 3-(4,5-dimethylthiazol-2-yl)-2,5-diphenyl tetra sodium bromide(MTT) assay^28^. Cells were seeded in a 96 well plates with a cell density of 2500 cells in each well. Cells were treated with test drug, dissolved in DMSO (0.5%final conc.)for 48 h. Cells were then treated with MTT reagent for 4 h followed by the addition of DMSO to lyse the cells and solubilize the formazan crystals. The samples were read using an ELISA plate reader at a wavelength of 570nm. The amount of color produced was directly proportional to the number of viable cells. Inhibition of cell growth, in the presence of test compounds, with respect to the control wells was calculated as percentage inhibition.

#### 2.2.2. MK2 enzyme based assay

MK2 enzyme assay was optimized using HTRF KinEASE STKS1 kit from Cisbio. Briefly, the reaction was initiated by adding 1.91 ng recombinant human MK2 enzyme to STKS1 substrate-biotin in the presence of and 188nM ATP in a 384 well plate, in kinase assay buffer (20mM HEPES buffer, 5mM MgCl_2_ and 1mM DTT, pH 7.4) in the absence or presence of different concentrations of the compound(PF-3644022). Total volume of the reaction was 20μl. Test compounds were dissolved in DMSO and the final concentration of DMSO in the reaction was 0.1 %. After 30 min of incubation at room temperature, 10 µl of detection reagent containing STK-Antibody, labeled with Eu3+-Cryptate and streptavidin-XL665 and EDTA(proprietary compositions of Cisbio) was added to the reaction well. After 1h incubation at room temperature, fluorescence was measured at an excitation wavelength of 337nm and dual emission wavelengths of 665nm and 620nm in the Pherastar micro-plate reader(BMG Lab Technologies). Dose response curve was generated using different concentrations of inhibitor and IC_50_ was calculated using Graph Pad Prism.

#### 2.2.3. TNF-α induced CXCL1

A549 cells were plated at a density of 50,000 cells per well in a 96 well plate. Cells were treated next day with test compounds for 30 minutes followed by treatment with TNF-α at 5ng/ml for 16h. Cell supernatants were collected and levels of CXCL1 were evaluated by ELISA method.

#### 2.2.4. Inhibition of TNF-α, IL-6 and IL-10 in PBMCs induced with LPS by ELISA

Effect of test compounds on the levels of LPS induced release of cytokines was evaluated using human PBMCs. Briefly, hPBMCs were plated at a density of 1×10^5^ cells per well in a 96 well plate. Cells were pre-treated with the inhibitor for 30 min, followed by stimulation with LPS(1 μg/ml) and incubated further at 37°C in a CO_2_ incubator for 24h Cell culture supernatants were evaluated for TNF-α and IL-6 and IL-10 by ELISA.

#### 2.2.5 LPS induced phosphorylation of HSP27and JNK1/2 in U937 cells by ELISA

U937 cells were plated at a density of 2.5x 10^5^ cells per well in a 24 well plate. Cells were pre-treated with test compound for 1h, followed by stimulation with LPS(100ng/ml) for 60 min. Cell lysates were prepared and analyzed for phospho-HSP27(Ser78) or phospho JNK1/2(Thr183/Tyr185) by ELISA using DuoSet ELISA kits(R & D Systems) as per the manufacturer’s instructions. Test compounds were dissolved in DMSO and final concentration of DMSO in all the cell based assays was 0.5 %.

#### 2.2.6 Phosphorylation of LPS induced p38 MAPK, JNK and Msk1/2 in U937 cells by western blot

U937 cells were plated at a density of 1 × 10^6^ cells per well in a 6-well plate followed by overnight serum starvation. Cells were then pre-treated with the test compound for 60 min, followed by stimulation LPS(100ng/ml) stimulation for 30 min. Cell lysates were prepared and analyzed by western blot to check the levels of p-p38 MAPK and pMSK1/2 by western blot.

#### 2.2.7. LPS induced TNF-α mouse endotoxemia model

PF-3644022 was evaluated in LPS induced endotoxemia model as per to confirm its efficacy in animal model. Briefly, PF compound was administered by oral route, followed by LPS challenge at a dose of 25 μg/mouse by intranasal route. Animals were sacrificed after 4 hours and BAL analysis was performed and levels of TNF-α measured. Roflumilast was used as a method control in this study.

### 2.3 Statistical Analysis

Results were expressed as mean+ standard error of the mean of the three independent experiments Statistical analysis was done using one way univariate analysis of variance followed by Bonferroni’s test for multiple comparisons and Student’s t-test for single comparison.

## 3.0 Results

Our data gives us a very clear understanding of the mechanistic differences at the molecular level as a result of inhibiting two members of the same pathway under the similar experimental conditions. Our data confirms that although inhibiting both p38 MAPK as well as MK2 can be potential anti-inflammatory approaches, inhibiting MK2 can certainly be a better approach with regard to the potential toxicities associated with them.

### a) Mode of action of PF-3644022 and BIRB-96

PF-3644022 and BIRB-96 were first evaluated in *in vitro* assays to confirm their modes of action. Both the molecules were evaluated for their ability to inhibit the phosphorylation of p38 MAPK in PMA differentiated U937 cells after LPS challenge, using western blot. LPS stimulation led to the activation of phospho-p38 MAPK protein in U937 cells as compared to unstimulated cells. Pre-incubation of 10 and 1μM of BIRB-796, resulted in the significant reduction in the levels of LPS-induced phospho-p38 MAPK. PF compound on the other hand did not show any inhibition of phospho-p38 MAPK(Figure 1a).

**Figure 1:**
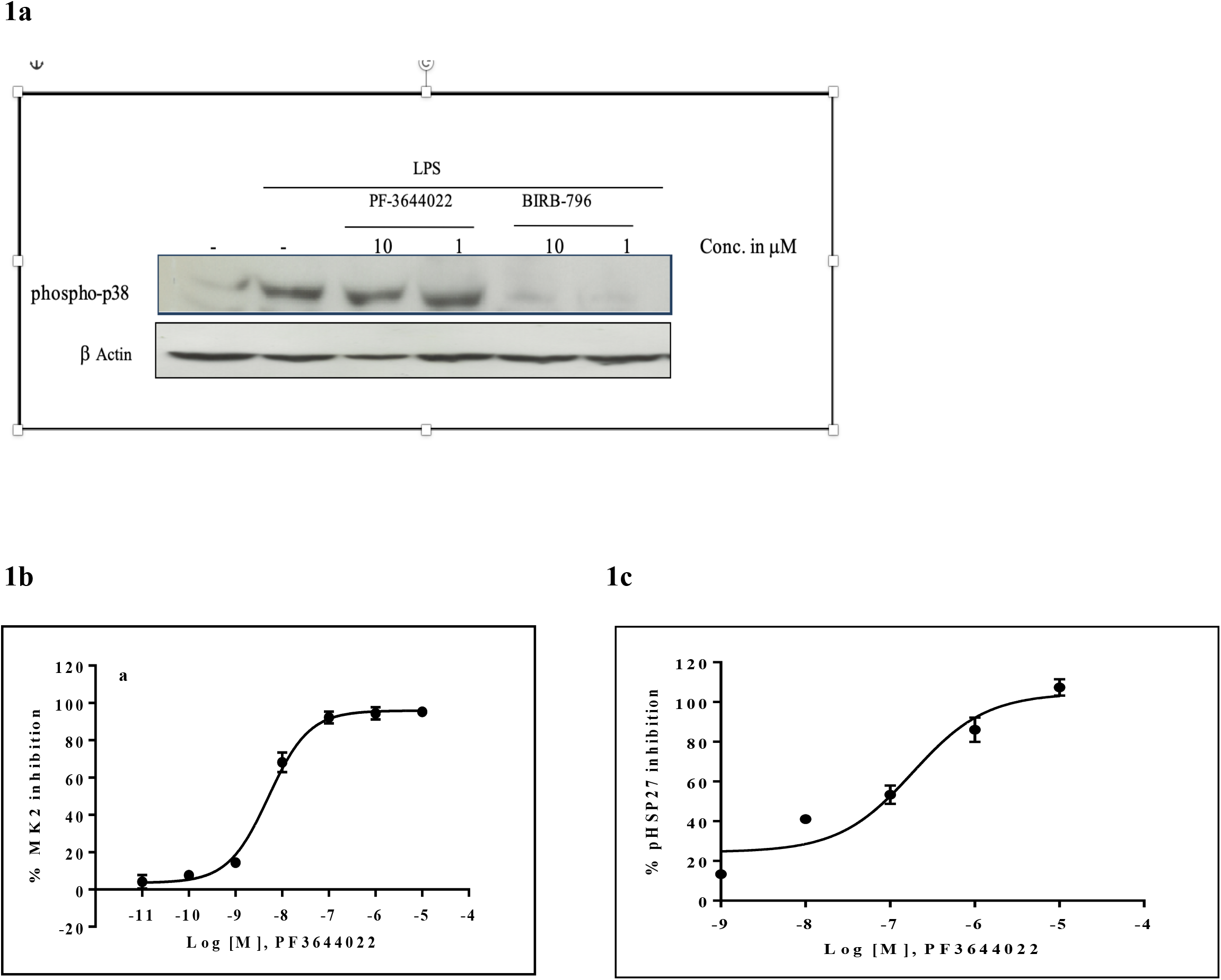
a) LPS-induced p38-phosphorylation in U937 cells; b)inhibition of MK-2 enzyme activity by PF-3644022 using KinEASE STKS1 method; c) inhibition of LPS-induced HSP27-phosphorylation in U937 cells by PF-3644022

Further to this, PF compound was evaluated in cell free MK2 kinase assay using HTRF KinEASE STKS1 kit. MK2 enzyme was titrated to determine its optimum concentration. ED_80_ values of the enzyme and Km of ATP were determined prior to the inhibition assay and found to be 1.91ng and 188nM respectively(Data not shown). IC_50_ value of PF-3644022 for MK2 enzyme inhibition at EC_80_ concentration of enzyme and ATP, at its K_m_ value was found to be 5.18nM(Figure 1b).

Since Hsp-27 is a direct substrate of MK2, PF-3644022 was also evaluated to confirm its ability to inhibit LPS induced phosphorylation of Hsp-27 in U937 cells and its IC_50_ value was found to be 86.4nM by ELISA method(Figure 1c).

### b) Anti-inflammatory properties of p38 and MK2 inhibitors

#### i) Inhibition of TNF-α induced CXCL1

Many p-38 MAPK inhibitors have progressed to phase II and III based on their potent anti-inflammatory profile for asthma and COPD. TNF-α is the predominant inflammatory stress in inflammatory conditions which is also responsible for neutrophil infiltration at the site of inflammation during bacterial and viral pulmonary exacerbation. CXCL1 is a key regular of neutrophil homeostasis. CXCL1 is remarkably expressed by macrophages, neutrophils and epithelial cells and has neutrophil chemoattractant activity. Increased concentrations of CXCl1 are also found in highly invasive fibroblast like synoviocytes(FLS) in rheumatoid arthritis patients. TNF-a antibody has been previously shown to inhibit TNF-a induced induction of CXCL1 in cell based models and animal studies. In our cell based studies using lung epithelial cell line A549, we have shown that TNF-α leads to induction of CXC1 and both p38 and MK2 inhibitors reduced the levels of TNF-α induced CXCL1(Figure 2).

**Figure 2.**
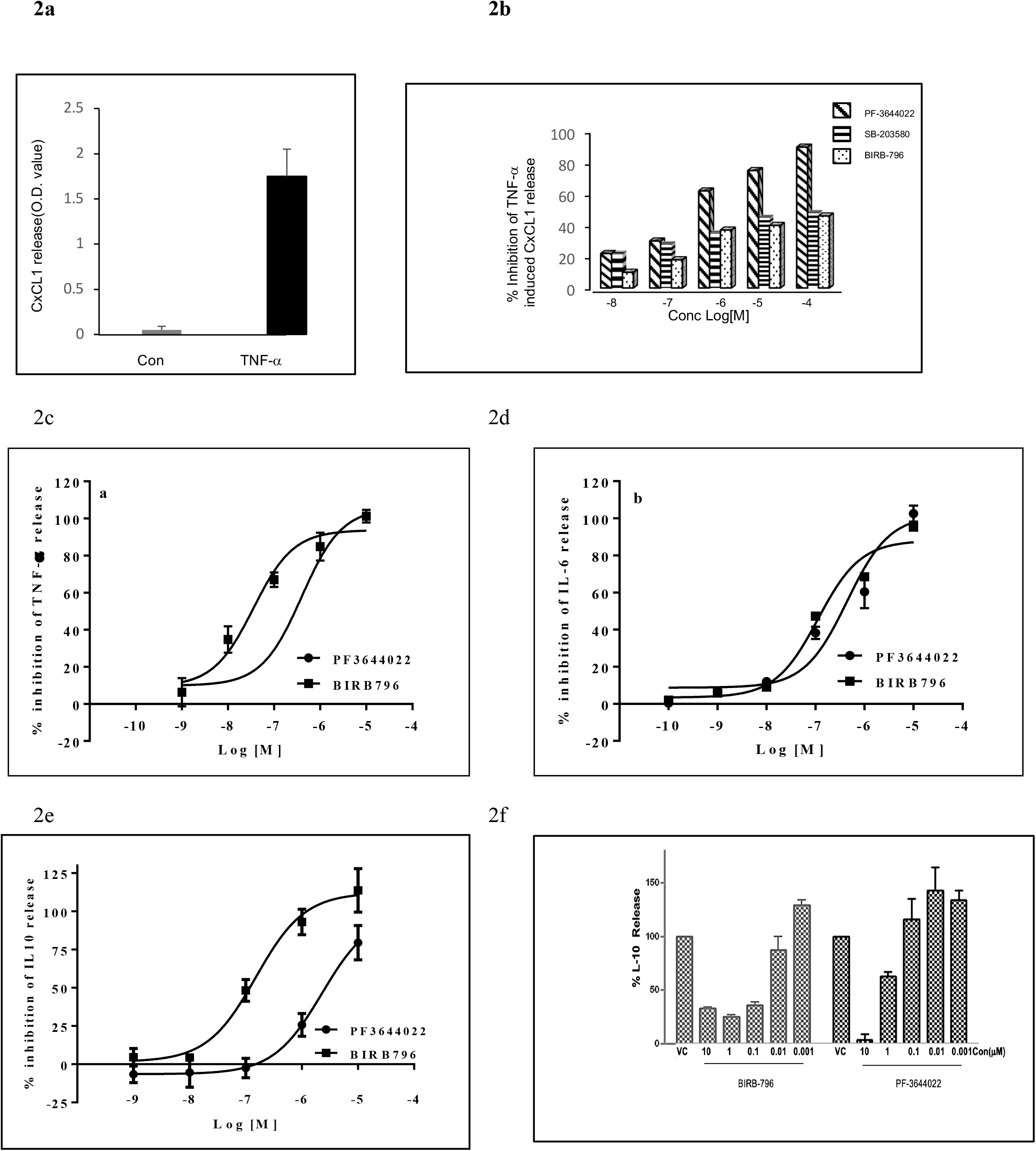
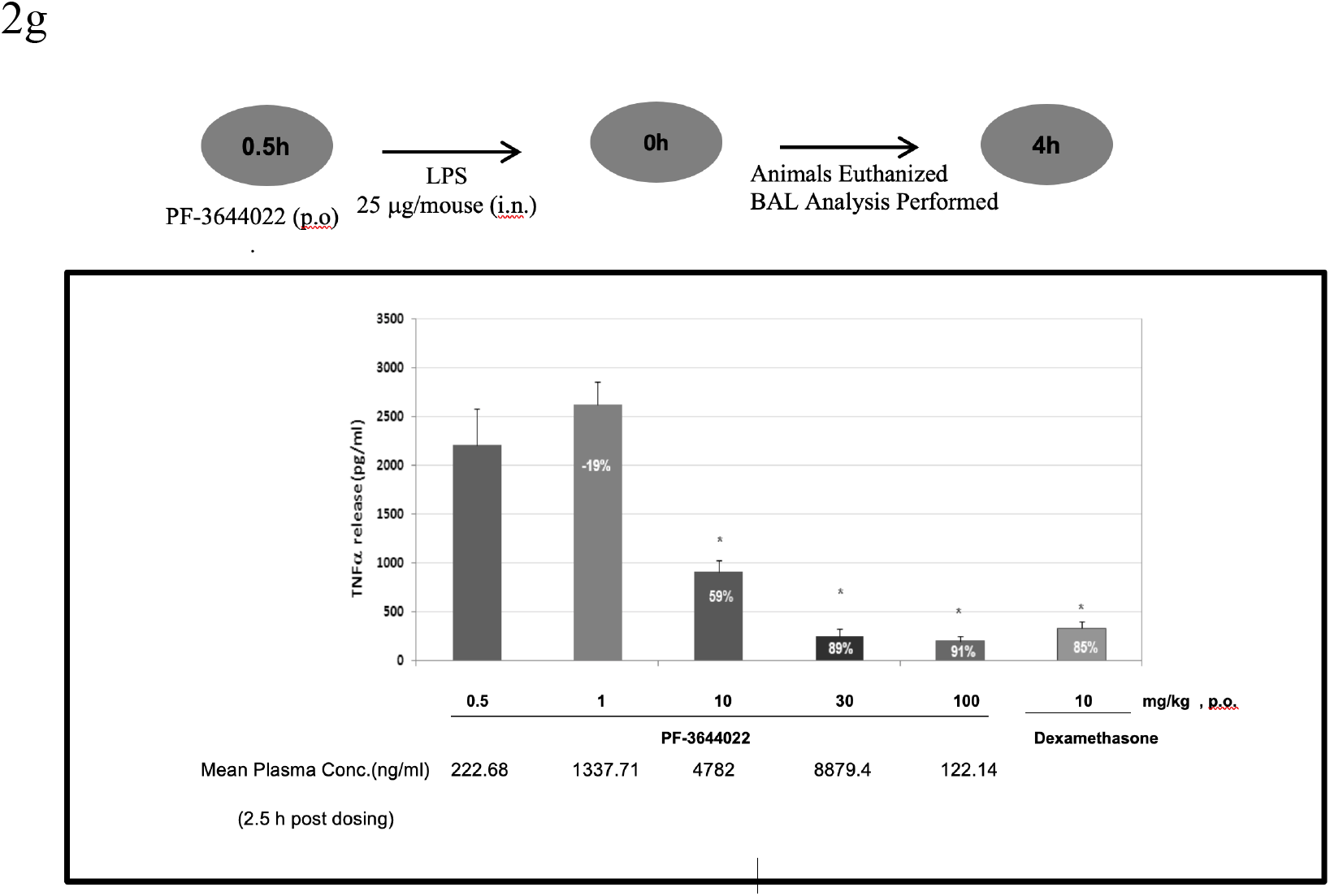
Anti-inflammatory potential of p-38 and MK2 inhibitors a&b) TNF-α induced release of cxcl-1 in A549 cells and its inhibition by MK2 and p-38 inhibitors, c&d)inhibition of LPS induced TNF-α and IL-6 respectively in U937 cells by ELISA, e&f) inhibition of LPS induced IL-10 in human PBMCs by ELISA; g)efficacy of PF-3644022 in LPS induced TNF-α release mouse endotoxemia model using balb-c mice

#### ii) Inhibition of LPS induced TNF-α and IL-6 using human PBMCs

Effect of MK2 inhibitor, PF-3644022 and p38 MAPK inhibitor, BIRB-796 were evaluated for the release of TNF-α and IL-6 from LPS stimulated PBMCs. Both the inhibitors showed dose dependent inhibition of TNF-α and IL-6 using cell culture supernatants of LPS stimulated PBMCs(Figure 2, Table-1).

#### iii) Inhibition of IL-10

Because of the involvement of both pro and anti-inflammatory loops in p38 MAPK pathway, we also evaluated the likelihood of inhibition of LPS induced IL-10 production which is an anti-inflammatory cytokine by the p-38 MAPK and MK2 inhibitors(Figure 3). We found that p38 MAPK inhibitor, BIRB-796 potently inhibited the LPS-mediated induction of anti-inflammatory cytokine IL-10 secretion with an IC_50_: 118nM, whereas, PF-3644022 showed very weak inhibition of IL-10 with an IC_50_ value of 2630nM in this experiment.

**Figure 3:**
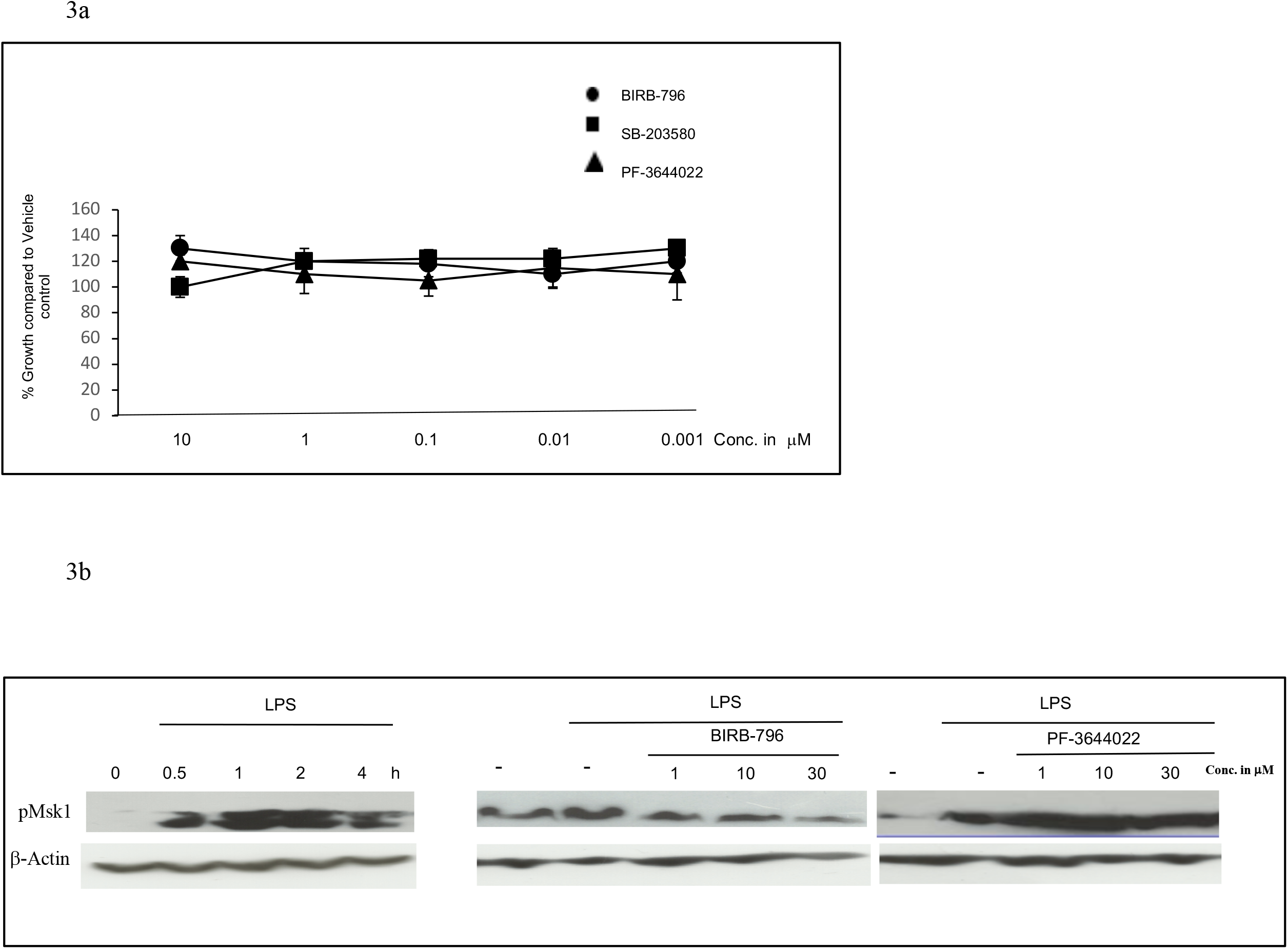
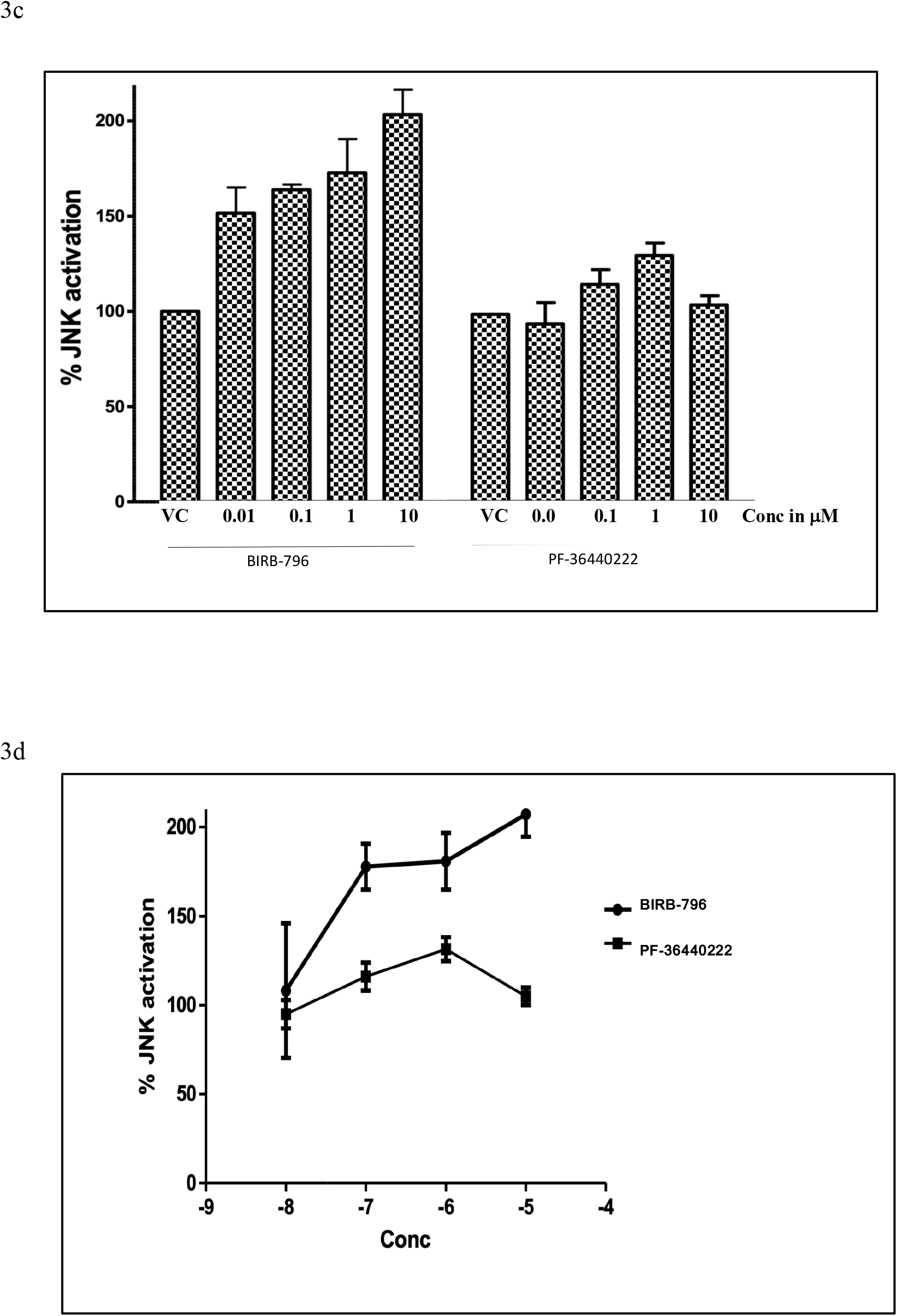
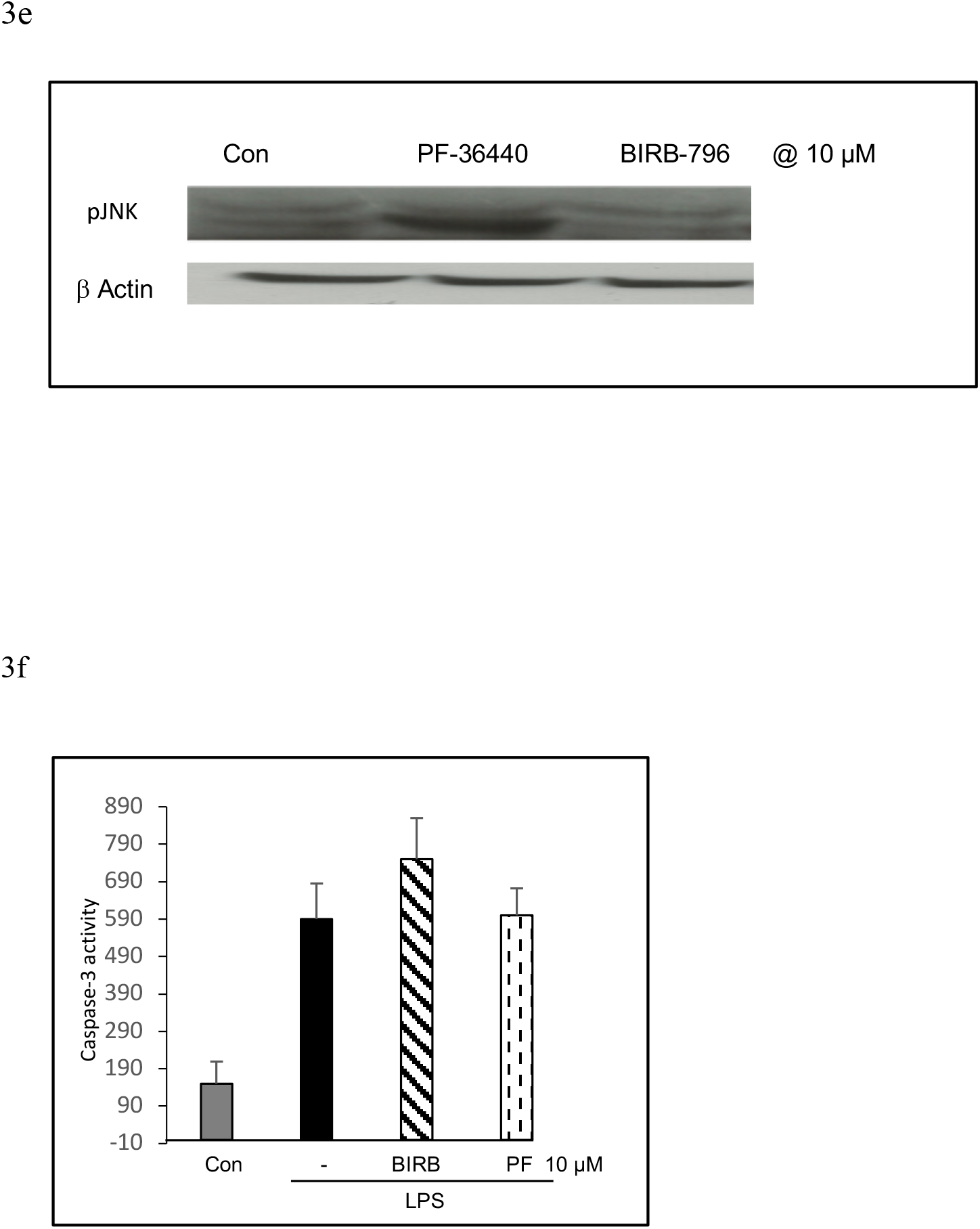
Assessment of toxicities by p-38 MAPKK and MK2 inhibitors in cell based assays a)Cell proliferation assays in A549 cells by MTT method, b)LPS induced pMSK1/2 in U937 cells by western blot; c&d) JNK activation in LPS-induced U937 cells by ELISA method and e) by western blot, f) caspase-3 release in LPS induced U937 cells by ELISA method.

#### iv) Efficacy in mouse endotoxemia model

To confirm the *in vivo* efficacy, PF-3644022 was evaluated in LPS induced endotoxemia model. PF-3644022 showed dose related inhibition of TNF-α release when administered by oral route. Compared to LPS control and ED_50_ value of PF-3644022 was found to be 7.02 mg/kg in this model. Roflumilast was used as a method control in this study and demonstrated 85 % inhibition of LPS induced TNF-α at 10 mg/kg by oral route(Figure 1d).

### b) Cellular toxicities of p38 and MK2 inhibitors

#### i) Cell proliferation assay using MTT assay

p38 MAPK inhibitors SB-203580, BIRB-796 and MK2 inhibitor PF-3644022 were evaluated in the cell proliferation assay using MTT but none of these molecules showed any toxicity in this assay(Figure 3). This data proposed the need to develop other cell based assays which can be used as test models to demonstrate the reported toxicities of p38 MAPK inhibitors in clinic and also as a tool to predict the toxicities of proposed MK2 inhibitors or other anti-inflammatory molecules in the pipe line.

#### ii) Inhibition of LPS induced pMsk1/2 in U937 cells

PF-3644022 and BIRB-796 were evaluated for the inhibition of LPS induced Msk1 phosphorylation in PMA differentiated U937 cells. LPS challenge resulted in the phosphorylation of Msk1 protein as compared to un-stimulated cells in western blot study. p38 MAPK inhibitor in this experiment led to inhibition of LPS induced Msk1/2 phosphorylation as compared to LPS-stimulated cells. However, MK2 inhibitor, PF-3644022 did not show any inhibition of LPS induced phosphorylation of Msk1 in these cells up to 30 μM(Figure 3).

#### iii) Activation of p-JNK in U937 cells

Effect of p38 MAPK inhibitor BIRB-796 and MK2 inhibitor PF-3644022 were evaluated for JNK activation in U937 cells, induced with LPS. LPS challenge resulted in phosphorylation and therefore activation of JNK protein as compared to un-stimulated cells. p38 inhibitor, BIRB-796 although led to the significant activation of JNK protein in LPS-treated cells as compared to LPS-stimulated cells, MK2 inhibitor, PF-3644022 however did not show activation of p-JNK levels up to 10 μM as evaluated by ELISA as well as western blot(Figure 3).

## 4.0 Discussion

Progression of several p-38-MAPK inhibitors from early discovery to late stage clinical evaluation has already provided a proof of concept for p38 MAPK inhibitors having strong anti-inflammatory response, although with genuine concerns about their transient efficacy and adverse toxicity. MK2 inhibitors on the other hand, are at the stage of infancy. In this study, we generated basic *in vitro* and *in vivo* data to demonstrate the anti-inflammatory potential of p38 MAPK and MK2 inhibitors. Further, we conducted several cell based assays to investigated the mechanistic differences of inhibiting the p38 MAPK and its downstream substrate MK2 by. p38 MAPK and MK2 inhibitors used in this study were BIRB-796 and PF-3644022 respectively. BIRB-796 is one of the most potent^10,11^allosteric inhibitor of p38-α isoform and binds with slow association and dissociation rates^12^. Some basic data is also available suggesting the efficacy of MK2 inhibitors in both acute and chronic models of inflammation^13,14^ and inflammation associated colorectal cancer^15^. However, the volume of data is much less in comparison to the former. In the present manuscript, we have conducted a series of experiments to dig out the real advantages, if any of one over the other.

*In vitro* cell based data generated at our end clearly demonstrates that both p38 MAPK and MK2 inhibitors have strong anti-inflammatory properties by potently inhibiting the levels of LPS-induced a TNF-α and IL-6 in hPBMCs. However, p38 MAPK but not MK2, phosphorylates Msk1/2 which in turn increases the production of anti-inflammatory cytokine, IL-10. We have shown that BIRB-796 led to a significant reduction in the phosphorylation of pMsk1/2 as well as the levels of IL-10 in LPS-induced hPBMCs while PF-3644022 did not show such an effect very significantly. This can probably be one of the reasons, p38 MAPK inhibitors demonstrate a disbalance of pro- and anti-inflammatory properties. MK2 being downstream of p38 MAPK, spares Msk1/2 and therefore MK2 inhibitors do not inhibit pMsk1/2 or IL-10 and therefore they are not expected to cause any disbalance in their anti-inflammatory potential.

In addition to above, our data suggests that BIRB-796 activates p-JNK activity in LPS stimulated cells *in vitro*. PF-3644022, however, didn’t show any significant increase in the levels of p-JNK. Increased levels of p-JNK by p38 MAPK inhibitors have already been reported to cause liver toxicity, tumor growth^16,17^ and several other side effects associated with them^-19^. However, MK2 by not participating in the feedback signaling loop, MK2 inhibitors do not lead to the activation of JNK and therefore are expected to be less toxic^20^.

Based on the evidences obtained from our comparative cell based mechanistic assays, we believe that inhibition of p38 MAPK may be associated with undesirable side effects due to its association with several redundant signaling partners. Inhibiting MK2 on the other hand can be a more specific and relevant target for anti-inflammatory therapy(Figure 4). Although, first generation, ATP-competitive MK2 inhibitors have suffered from low solubility, poor cell permeability, and scarce kinase selectivity, new non-ATP-competitive inhibitors of MK2 are being developed to circumvent the selectivity issues^25^. MMI-0100, an MK2 inhibitor from cell-penetrating peptides(CPPs) has shown good safety and tolerability profile in three phase 1 clinical trials for idiopathic pulmonary fibrosis(IPF) as a dry powder inhalant^26^. Targeting MK2 therefore suggests a promising therapeutic opportunity for inflammatory disorders.

**Figure 4:**
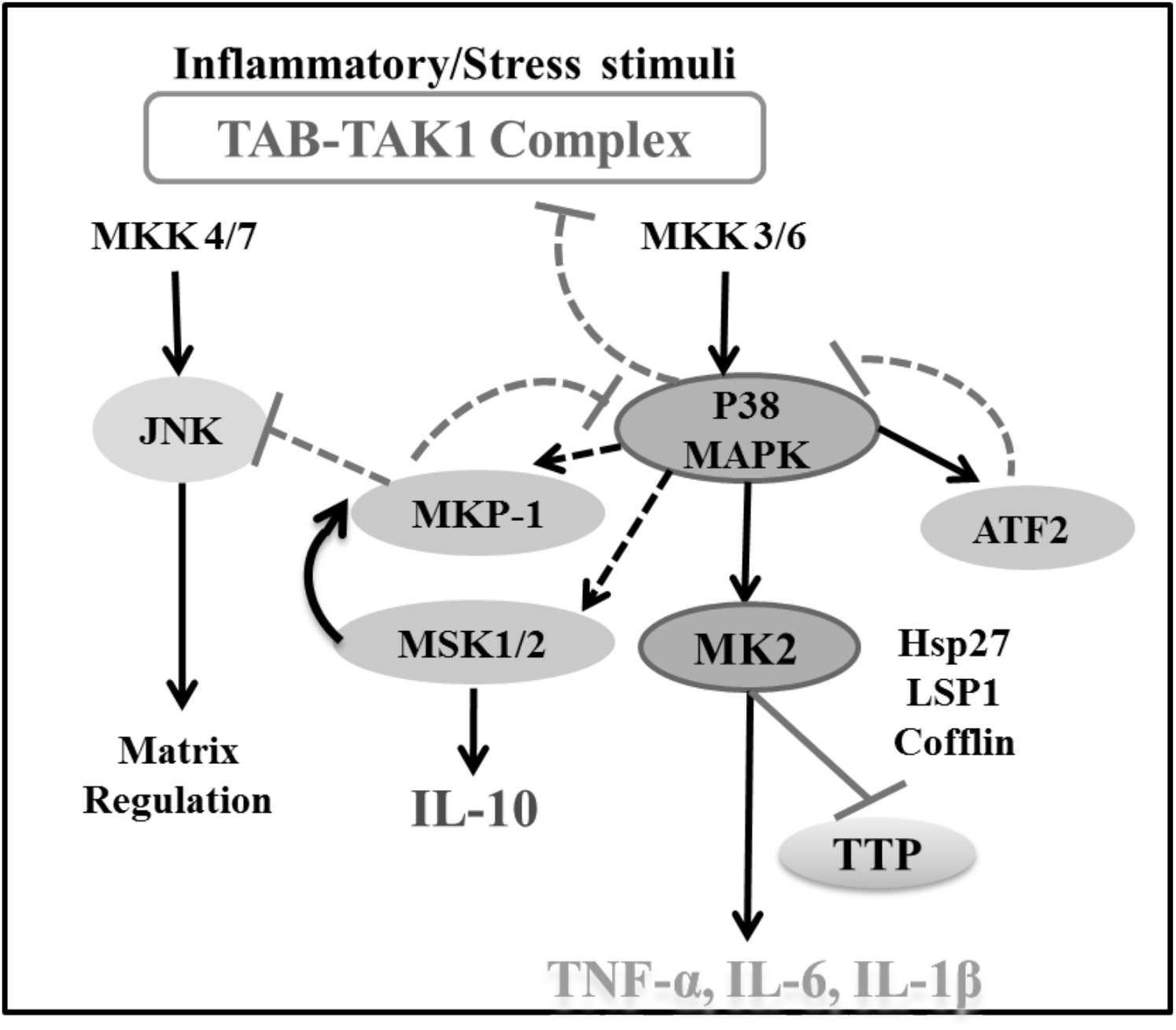
Schematic representation of p38MAPK pathway. p38 is involved in the regulation of pro-inflammatory (cytokines), and anti-inflammatory (inhibition of TAK1 and JNKs, regulation of MKP-1, MSK1/2 and IL-10) loop. The downstream MK2 is only involved in the production of pro-inflammatory cytokines.

## Notes

### Competing Interest Statement

The authors have declared no competing interest.

